# Analysis of Gene Expression Differences Between Eastern and Western Loblolly Pine Seed Sources

**DOI:** 10.1101/2023.04.24.538124

**Authors:** Adam R Festa, Ross Whetten

## Abstract

The selection of an appropriate seed source for a given geographic region is critical to ensuring prosperous southern pine plantations. The observed variation between eastern and western loblolly pine seed sources has shown differences in economically advantageous traits such as drought tolerance, growth rates, and disease resistance. Understanding what drives these local adaptations is of interest, given that current forecasted climate modeling suggests there will be increased temperatures and changes to precipitation by the year 2050. The objectives of this experiment were to 1) identify differentially expressed transcripts between eastern and western loblolly pine sources; 2) link these transcripts to *Arabidopsis* orthologs; 3) compare GO categories of differentially-expressed transcripts. The findings highlighted include interesting pathways and genes that are related to the known differences among eastern and western seed provenances. Additionally, they represent fundamental differences in the beginning of seedling development without any treatment or disease pressure applied, showing that there are detectable differences between these two provenances at a young age. Overall, this experiment contributes to the body of literature on fundamental differences between loblolly pine seed sources.

## Introduction

Global temperatures are expected to rise two degrees Celsius by 2050 (Yerlikaya et al. 2020). Alongside this temperature rise is the expectation of increased global carbon emissions and variability in local precipitation rates (Panagos et al. 2022, OECD 2012). These climate changes are expected to affect agricultural and forestry productivity (Beach et al. 2015, Howden et al. 2017, Malhi et al. 2021). While the impact is predicted to decrease agricultural productivity, the increase in CO_2_ has been proposed to initially increase forest productivity (Kirilenko and Sedjo 2007, Arora 2019, Cammarano et al. 2022). However, it is not clear how the changes in weather patterns will affect incidence of drought, flooding, or disease pressure. Current efforts on adaptation strategies developed to mitigate the impact of climate change will presumably have to be deployed on a regional or local basis (Lambert et al. 2021). How agriculture or forestry breeders will need to adapt depends on a wide range of factors such as their natural geographic range, available genetic variation/resources, and economic impact.

*Pinus taeda* L. (loblolly pine), the most planted tree across the Southern US with just over 400 thousand hectares planted annually, has a widespread natural range from the North to Maryland, South to Florida, and West to Texas in the United States (McKeand et al. 2020, Schultz 1997). Across this geographic range, there has been extensive evidence of physiological and phenotypic differences among seed sources (Szymanksi and Tauer 1991, Jayawickrama et al. 1998, Sierra-Lucero et al. 2002). The general patterns observed include differences in drought tolerance, resistance to fusiform rust (*Cronartium quercuum* f. sp. *Fusiforme*), and overall growth. Loblolly pine families from west of the Mississippi River are known for slow growth, drought tolerance, and disease resistance. In contrast, families from east of the Mississippi River typically display fast growth and have moderate disease resistance but are prone to drought stress. A recent review article produced by Matallana-Ramirez et al. (2021) provides an excellent overview of the potential impacts of climate change on loblolly pine and the possibility of breeding for adaptation.

With respect to the effect of temperature increase on growth, research is limited, and results are a factor of seed sources tested and environmental variables assumed. For example, Schimdtling (1994) utilized historical provenance test data to predict the impact of a 4.0°C temperature increase on height and found a general 5-10% reduction. In another study, Nedlo et al. (2009) evaluated biomass accumulation on 34 seedlings from a single open-pollinated Georgia family planted across four sites ranging from North Carolina to the middle of Florida, representing an 8.7°C difference in mean growing season temperature. The evaluation of biomass accumulation was highly correlated with the absolute difference in mean growing season temperature and was reported being 48% less when planted North and 43% less when planted South.

Similarly, there are few predictions of how loblolly pine families may respond to climate change when including both temperature increase and precipitation variability. Farjat et al. (2015) modeled the potential effects on height growth for eastern loblolly pine sources given a 5% decrease in precipitation and 0.5°C change in max or minimum temperatures. They reported that pine families in Georgia, Florida, and South Carolina showed an increase in performance relative to current environments; however, they declined in height as they were moved north or inland. Another study obtained exome-wide genotype data for 299 trees from east of the Mississippi River, ranging from Delaware to Florida, and predicted the adaptive capacity of populations leveraging associations between SNPs and current climate variables versus projected allele frequencies given future climate variables. They found that tree populations in the northeastern part of the natural range, including Delaware, Maryland, Virginia, North Carolina, South Carolina, and Northern Georgia, were most likely to be impacted (Lu et al. 2019).

The local adaptation of loblolly pine families and how they may respond to changes in weather patterns are inherently dependent on underlying genetic differences. Beyond using SNP markers derived from DNA, another way of understanding the fundamental genetic differences in seed sources may be accomplished using gene expression. For example, the use of a microarray analysis containing 2171 expressed sequence tags collected from loblolly pine xylem and shoot tip from two seed sources, South Arkansas and South Louisiana, were grown on a common site and found that variation in gene expression may reflect local adaptation (Yang and Loopstra 2005).

In addition to understanding local adaption, gene expression has been used to evaluate transcripts or genes that may be involved in response to abiotic stresses such as induced drought. One example in loblolly pine subjected three unrelated clones to three different water regimes. The three clones were derived from the eastern provenance. After applying water treatments, the researchers’ generated cDNA libraries from root tissue. The resulting analyses identified significant transcripts previously associated with drought tolerance in other species. They also reported finding most of these transcripts in one clone, highlighting that allelic variation and dependency on an individual’s genotype may be more important than treatment (Lorenz et al. 2006).

Over time, significant advancements have been made in sequencing technology, statistical approaches to identify differentially expressed genes, and the ability to leverage functional information (Han et al. 2015). The current standard approach for differential-expression analyses generally starts with RNA extraction, library creation, and sequencing. Raw reads returned from sequencing are then mapped against a reference genome and utilized in one of the various differential expression algorithms to test for significance.

There are a variety of challenges and opportunities for systematic bias to be introduced when conducting tests of significance for differential expression (Finotello and Di Camillo 2015). For example, when comparing two samples, different sequencing depths can affect comparisons and lead to claiming something is differentially expressed when expression differences are due to lack of coverage. Additionally, different gene lengths and GC content have been shown to produce bias even within a single sample. These are all essential factors to take into consideration when determining significance.

Once differentially expressed transcripts are identified, more practical information about these genes can be found using gene ontology (GO) or Kyoto Encylopedia of Genes and Genomes (KEGG) (The Gene Ontology Consortium 2019, Kanehisa et al. 2021). Gene ontology is a public resource that contains biologically relevant information about known genes and proteins. The database is structured into three categories: cellular components (CC), molecular functions (MF), and biological processes (BP). The enrichment of GO categories is typically assessed using statistical analyses of GO categories within the up- and down-regulated transcripts found to be differentially expressed in order to identify over- or underrepresented terms. Like the dependency of a gene being differentially expressed biased by its length, the likelihood of a GO category found is affected by the length and number of genes within a category, so it should be considered when determining significance (Young et al. 2010).

The KEGG database contains three categories: pathways, BRITE and Modules. Genes found to be differentially expressed are typically mapped to the KEGG database to further understand biologically relevant metabolic pathways they may impact. The Modules category contains specific metabolic capabilities and those gene sets involved, and the BRITE categories group genes based on their function, such as enzymes or transcription factors.

RNA-Seq differential expression in conjunction with GO and KEGG pathways has been used extensively and provided beneficial insights to model species and row crops (Meng et al. 2022, Xie et al. 2019, Cao et al. 2021, Chen et al. 2021, Wang et al. 2021). However, both the GO and KEGG databases contain genes for which known functions have been studied. For species with little gene annotation, a common approach is to use BLAST to identify genes or protein sequences with similar potential orthologs to connect functional information to differentially expressed transcripts (Camacho et al. 2009, Zhang et al. 2012, Tian et al. 2022). Given the lack of available gene information for forest tree species, including loblolly pine, it makes sense that current differential expression experiments use BLAST to identify likely orthologs in model plant species (Heredia et al. 2016). A more recent loblolly pine study conducted a differential-expression analysis on RNA-sequencing of root tissue collected from three different genotypes under well-watered or low-watered treatment. Transcripts identified as being differentially expressed were queried against the NCBI BLAST database and performed GO enrichment as well as analysis of KEGG pathways. Significant changes in over 4,000 drought-related transcripts were reported, and gene ontology enrichment included categories such as response to water deprivation, and KEGG pathways revealed upregulation of Glutathione metabolism (Li et al. 2021).

Provided the wide geographic range, observed phenotypic variation among seed sources, and potential impact climate change may bring, the opportunity exists to further explore underlying genetic differences that may play a role in local adaptation for loblolly pine. Previous literature utilizing RNA-sequencing to explore these differences have been limited in their use of multiple families representing a wide geographic range. Additionally, the experiments have applied treatments in order to enhance changes in expression related to specific phenotypes. In the face of climate change, understanding what drives phenotypic variation observed from the wide geographic range of loblolly pine without applying treatments seems worthwhile to explore.

The objectives of this experiment were to utilize RNA sequencing on 6-week-old seedlings to evaluate any detectable differences in gene expression between eastern and western sources of loblolly pine families. More specifically, the aim is to 1) identify differentially expressed transcripts between multiple families representing the eastern and western loblolly pine sources; 2) link differentially expressed transcripts back to *Arabidopsis* orthologs; and 3) compare the enrichment of GO categories and KEGG pathways identified among the differentially expressed genes.

## Materials and Methods

### Experimental design

The plant material used in this experiment corresponds to batch 2 from Festa and Whetten (BioRxiv 2023, in review). That manuscript explains the methodology for germination, sample collection, RNA extraction, and sequencing. Of the 40 families grown in batch 2, 22 families were selected, comprising 11 Atlantic Coastal loblolly pine selections from North Carolina and South Carolina and 11 Western Gulf selections. The 11 eastern families had 29 biological replicates, and the 11 Western Gulf families had 32 biological replicates (Supplementary Table 1). Before analyzing differential gene expression, raw counts were transformed using the variance stabilizing transform function within *DESeq2*, and a principal component plot was generated using the function *plotPCA* (Love et al. 2014) and *ggpubr* R package (Kassambara 2020).

### Estimation of differential expression

The raw read count matrix containing 61 biological replicates was input to *DESeq2* version 1.30.1 (Love et al. 2014) to test differential expression between the western gulf and eastern loblolly pine families. All 32 biological replicates corresponding to the Western Gulf were contrasted against all 31 biological replicates representing the eastern families. DESeq2 implements a generalized linear model with a negative binomial distribution:

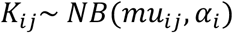

where counts *K*_*ij*_ for gene *i*, sample *j* are modeled using a Negative Binomial distribution with fitted mean *mu*_*ij*_ and a gene-specific dispersion parameter α_*i*_ The fitted mean can be defined as:

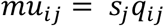

where *s*_*j*_ is the sample-specific size factor but can be substituted for gene-specific normalization factors and *q*_*ij*_ is a parameter related to the expected actual number of fragments for sample *j* .The parameter *q*_*ij*_ can be estimated as

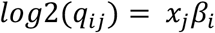

where the coefficients *β*_*ij*_ give the 2 *log*2 fold changes for gene *i* for each sample of the model matrix 𝓍.

To account for differences in count estimates given transcript length and GC content, instead of using sample-specific size factors, we replaced these with gene-specific normalization factors using the R package *cqn* (Hansen et al. 2012). The *cqn* package implements conditional quantile normalization for RNA-seq data. The resulting p-values of differentially expressed transcripts were corrected using the Benjamini and Hochberg method (Benjamini 1995), also known as the false discovery rate (FDR). Transcripts were considered differentially expressed if they had an FDR adjusted p-value less than 0.05. Results from differential expression were plotted using the R package *EnhancedVolcano* (Blighe et al. 2020).

### Annotation of differentially expressed transcripts

The functional annotation for each differentially expressed transcript was found by first querying against the complete *Arabidopsis thaliana* TAIR10 cDNA (Berardini et al. 2015) with NCBI command-line BLAST version 2.12.0 (Camacho et al. 2009) using tblastx and default parameters. The results of BLAST output were processed using the R package *rBLAST* (Hahsler et al. 2019). The top *Arabidopsis* cDNA hit for each loblolly pine transcript was kept, and only hits with an E-value less than 10^−5^ were retained as a possible match. *Arabidopsis* cDNA sequence names were converted to their corresponding Ensembl gene names (Cunningham et al. 2022), and gene ontology categories for each *Arabidopsis* gene were retrieved using the R package *biomaRt* (Durinck, 2009). The loblolly pine transcriptome contains multiple examples of several pine transcripts similar to a single *Arabidopsis* gene; we do not know if these similarities represent different alleles from loblolly pine or different members of gene families. In some cases, different pine transcripts in the up-regulated and down-regulated categories were similar to the same *Arabidopsis* gene. We don’t have enough information to interpret those results, so only *Arabidopsis* genes unique to either the up or down-regulated categories were kept.

The estimation of gene ontology categories that are over- or under-represented was accomplished with the R package *goseq* (Young et al. 2010). The *goseq* package, which helps overcome the selection bias of categories due to transcript length, utilizes a probability weighting function to estimate the likelihood of a gene being differentially expressed as a function of its length. The median length of cDNA for each *Arabidopsis* gene was estimated to define the null likelihood distribution of genes that would be differentially expressed. Resulting p-values were FDR adjusted and only categories with an adjusted p-value < 0.10 were retained. All plots were generated with *ggplot2* (Wickham 2016) and *ggpubr* (Kassambara 2020). To identify possible affected KEGG pathways, the unique *Arabidopsis* gene names in either the up or down categories were uploaded to the KEGG mapper (Kanehisa et al. 2021).

## Results

### Analysis of Illumina Sequencing Results

Across the 61 samples, total read counts per biological replicate ranged from 9.8 million reads to 39.2 million, with a mean of 25.0 million reads. For the 78,213 transcript contigs in the reference transcriptome assembly, read counts summed across all 61 biological replicates ranged from 0 reads to a maximum of 42.1 million, with a mean of 19,481 reads. After variance-stabilizing transformation, the PCA plot of read counts clustered into three groups: east, west, and far-west (Figure 1). The three families: LSG111, WG1809, and WG4801 clustered in the middle represent Western Gulf families selected from a geographic location between the east and west group samples (Figure 1).

**Figure 1.**
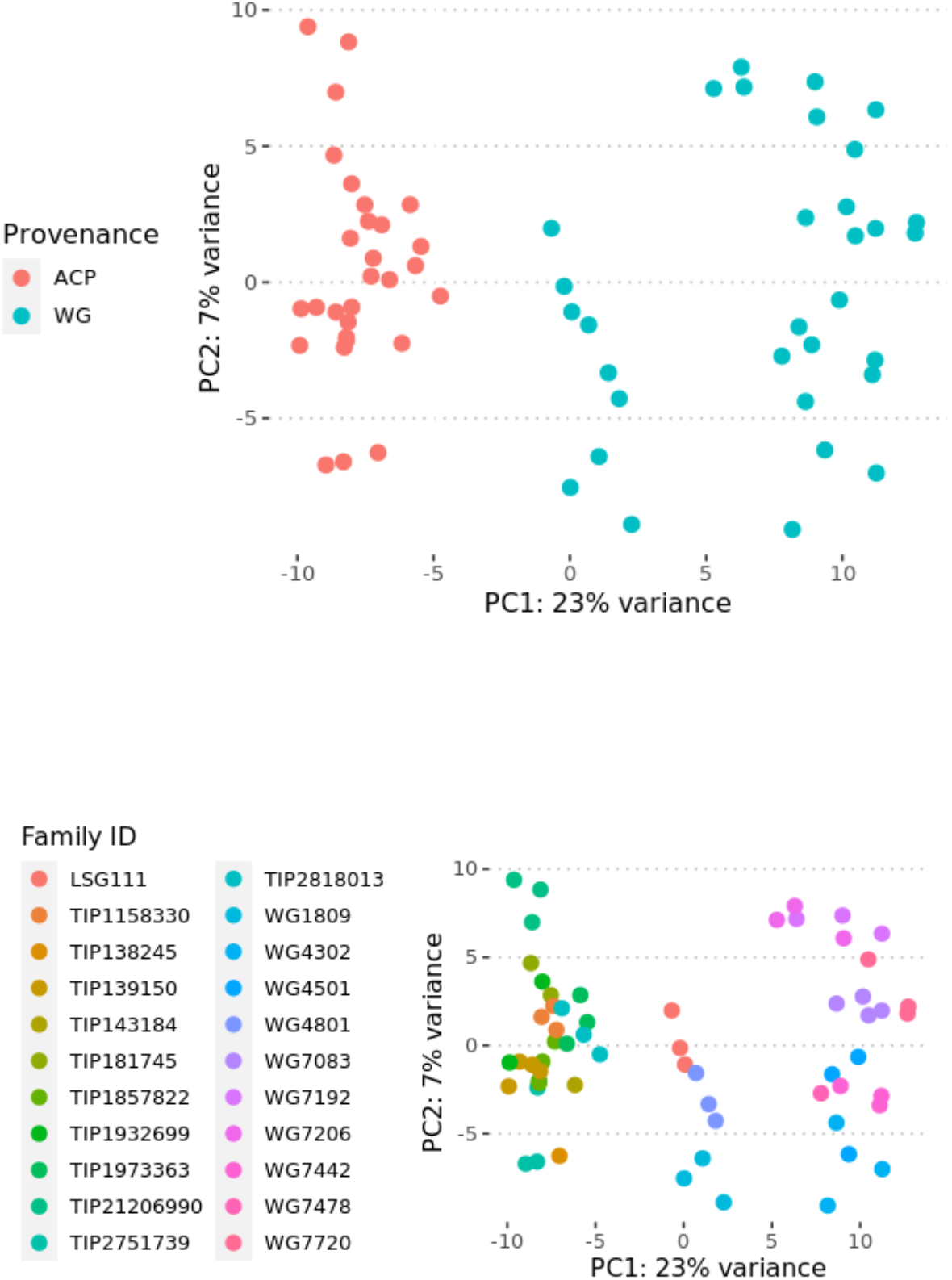
Principal component analysis of all 78,213 transcript gene expression estimates for biological replicates. (TOP) Biological replicates colored as their respective groups (N=29 ACP and N=32 WG), eastern families are red and western families are blue. (BOTTOM) Biological replicates colored according to their respective families (N=22). The eastern families are grouped on the left side of the plot, the far-western families are grouped on the right side of the plot, and three western families clustered in the middle (LSG111, WG1809, WG4801).

### BLAST results of Pita transcriptome to Arabidopsis

The 78,213 loblolly pine transcripts returned blast matches to 20,971 (43.4%) unique *Arabidopsis* cDNA sequences originating from 17,603 unique *Arabidopsis* genes. The distribution of mapping identities ranged from 14.29% to 100%, with a mean of 51.7%. Out of the 17,603 *Arabidopsis* genes, a total of 6911 genes were observed once, and the remaining were matched two times or more. After filtering blast results for matches with an E-value < 10^−5^, 57,996 Pita transcripts remained and matched 15,960 unique transcripts originating from 12,736 *Arabidopsis* genes. Mapping identities ranged from 14.69% to 100%, with a mean of 55.3%. The frequency of multiple pine transcripts matching a single *Arabidopsis* gene had a similar pattern in the filtered data to that observed across the whole data; 6554 *Arabidopsis* genes appeared once, and the remaining were found two or more times (Supplementary Table 2).

### Differential Expression between East and West seed sources

When comparing expression between the east and western loblolly pine biological replicates, approximately 4049 transcripts were upregulated in the western sources, and 5124 were downregulated (Figure 2). Within the transcripts found to be upregulated, 3,572 transcripts had hits to *Arabidopsis* genes with an E-value < 10^−5^. Of the 3,572 hits to *Arabidopsis* genes, 2304 unique *Arabidopsis* genes were found. A total of 1794 of these appeared once, and the remaining appeared two times or more. Within down-regulated transcripts, 4,567 hits had e-value less than 10^−5^, of which 2,396 were unique *Arabidopsis* genes. A total of 1748 out of 2396 were similar to a single pine transcript, and the remaining 648 were similar to two or more pine transcripts. Given the repetitiveness of matches, only the unique genes were kept. After combining the 2396 genes downregulated and 2304 upregulated genes, 3488 genes were unique, and 606 genes were in both categories, so they were removed from further analysis. Post filtering left 1790 unique *Arabidopsis* genes identified as downregulated and 1698 genes identified as upregulated. When evaluating functional annotation of differentially expressed pine transcripts, we utilize *Arabidopsis* gene names, based on the assumption that the pine transcripts are putative orthologs of the Arabidopsis genes.

**Figure 2.**
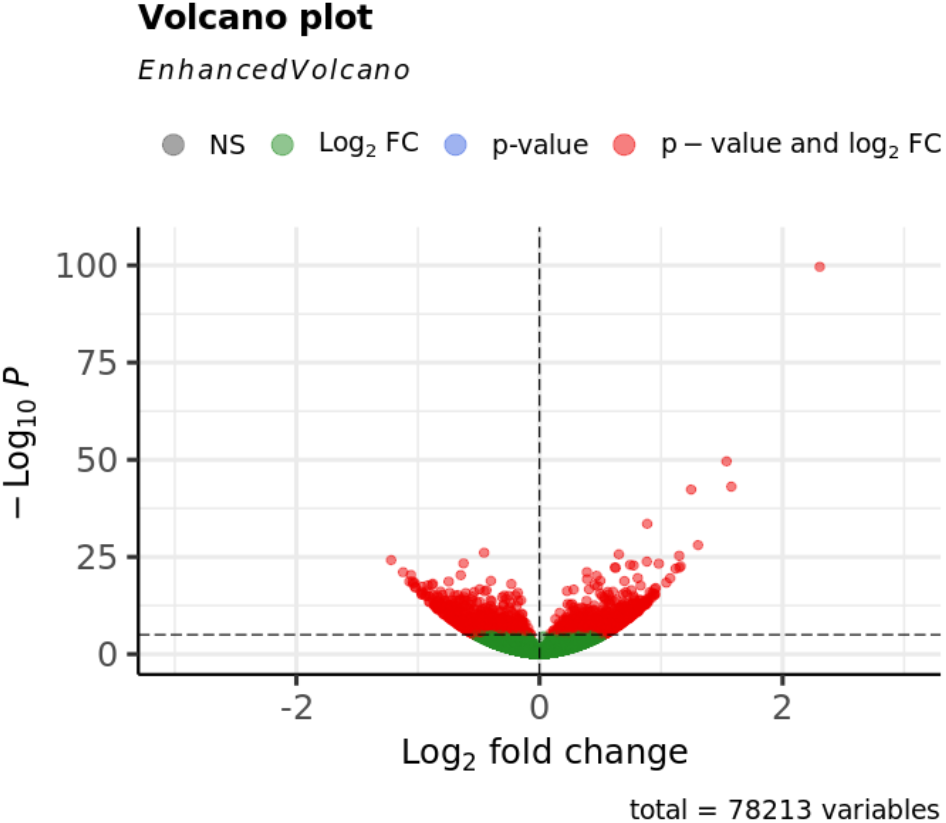
Volcano plot of differentially expressed transcripts between the western and eastern loblolly pine transcripts. A positive log2 fold change indicates higher expression in the western sources and a negative log2fold change indicates higher expression in the eastern pine transcripts. The -Log10P indicates the FDR adjusted significance of differentially expressed transcripts.

### Functional annotation of differentially expressed hits

After removing GO categories with a BH adjusted p-value < 0.1, downregulated *Arabidopsis* genes contained 72 categories: 20 biological processes, 25 cellular components, and 27 molecular functions. The upregulated GO categories that passed the BH threshold contained 114 categories: 59 biological processes, 32 cellular components, and 23 molecular functions. Only four overlapping terms were identified across both the upregulated and downregulated GO categories, including CC: cytosol and cytoplasm, BP: translation, and MF: mRNA binding.

We focused on the biological processes category because it is more informative to understanding physiological differences. The biological process GO terms enriched for upregulated pathways included: response to cold, response to heat, response to salt stress, response to water deprivation, and regulation of stomatal closure (Figure 3). The enrichment for downregulated biological process GO categories included defense response, protein phosphorylation, and sterol biosynthetic process (Figure 4).

**Figure 3.**
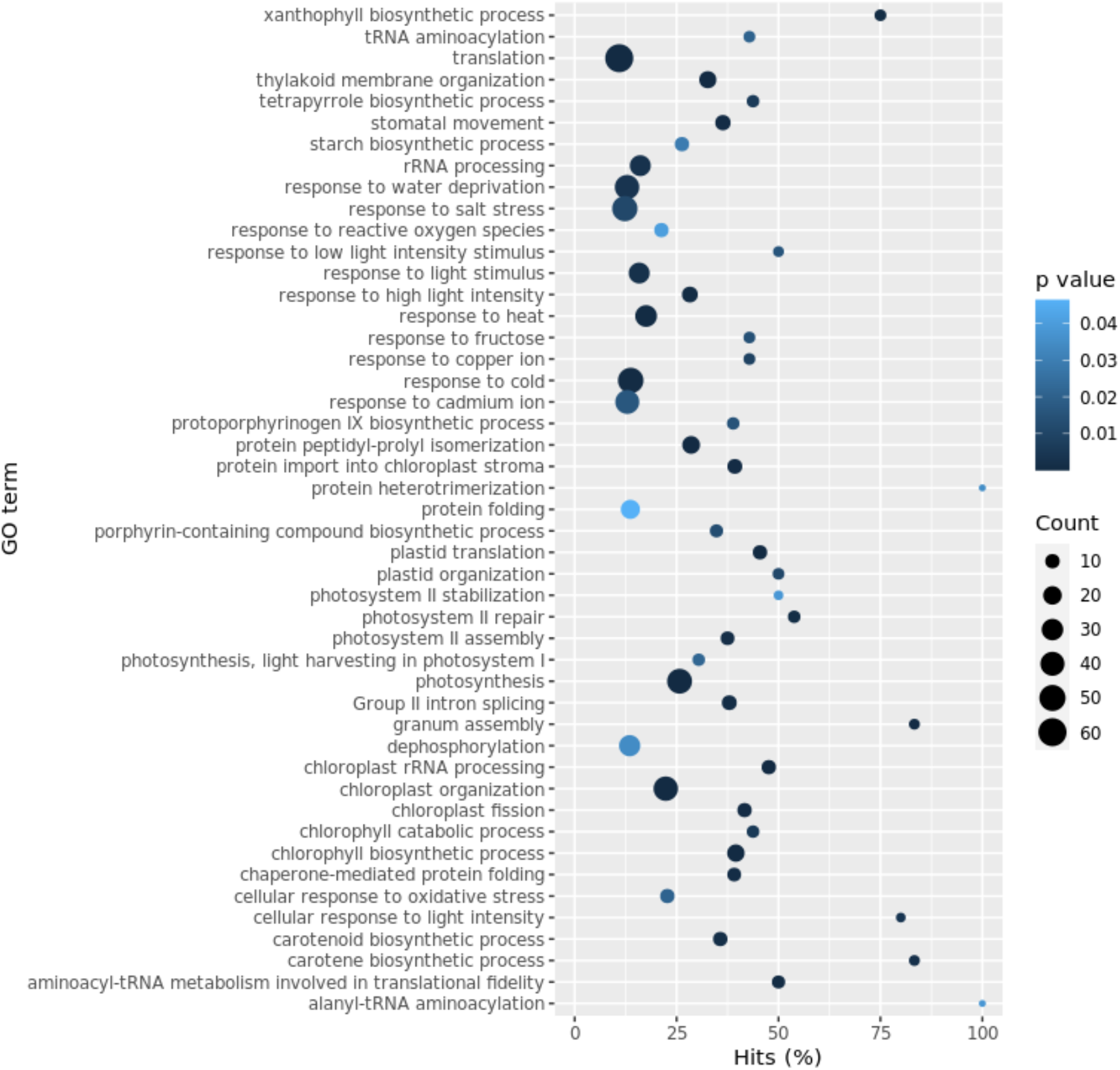
GO biological process terms with an FDR adjusted p-value < 0.05. overrepresented in the 1698 upregulated *Arabidopsis* genes. The color indicates adjusted p-value; size of the dot represents how many genes were identified; Hits % is the relative proportion of genes that were identified when compared to the total number of genes in that group.

**Figure 4.**
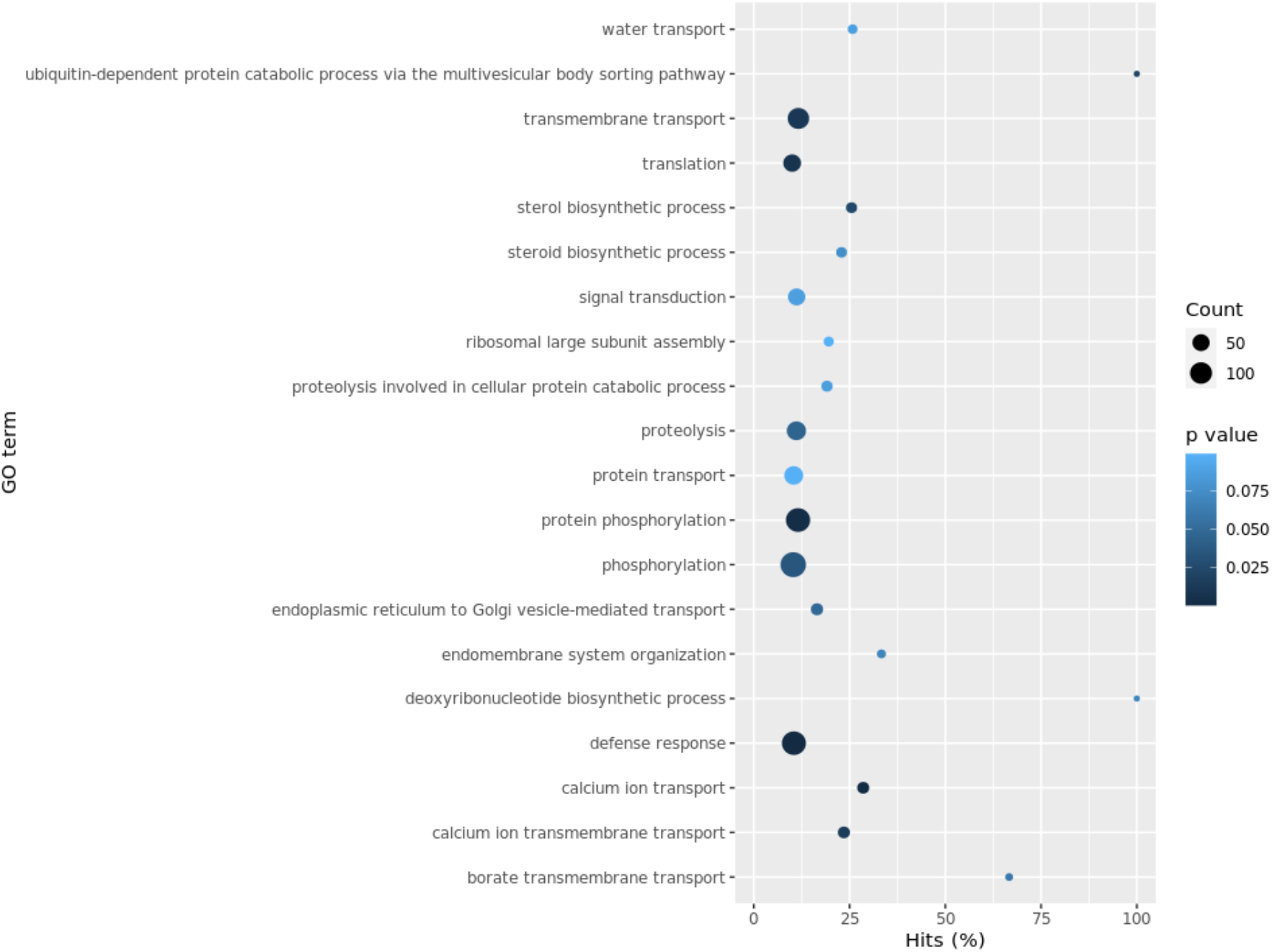
GO biological process terms with an FDR adjusted p-value < 0.10. overrepresented in the 1790 downregulated Arabidopsis genes. The size of the dot represents how many genes were identified; Hits % is the relative proportion of genes that were identified when compared to the total number of genes in that group.

### KEGG Pathway results of differentially expressed transcripts

The unique upregulated or downregulated *Arabidopsis* genes were uploaded to the KEGG mapper and resulted in each set having 123 pathways being affected. A total of 114 out of the 123 pathways overlapped and nine pathways were distinct to each set of *Arabidopsis* genes. The upregulated genes contained hits to 39 BRITE categories and 112 modules; the downregulated genes had hits to 41 BRITE categories and 97 modules. The KEGG web page (https://www.genome.jp/kegg/kegg3b.html) contains this definition of BRITE hierarchical categories: “The BRITE hierarchy file is created to represent functional hierarchy of KEGG objects identified by the KEGG Identifiers; for example, pathway-based gene classification or protein family classification by the K numbers.” Multiple pathways were found overlapping between both the upregulated and downregulated genes. For example, in the MAPK signaling pathway for salt and drought stress, HAB1, HAI3, ABI1, and HAI1 (all PP2Cs) and SnRK2.4 were upregulated, while PYL4, PYL7, and RCAR3 (all PYR/PYL) was downregulated. The plant hormone signal transduction pathway for zeatin biosynthesis returned RR16 (a type A-AAR) as being downregulated while RR1 (a type B-ARR) was upregulated. Within the BRITE category, several transcription factors linked to drought, salt, and stress tolerance were upregulated. A few examples of these included: TINY2, DREB2A, and bZIP2. Unique to the downregulated category was the impact of the brassinosteroid pathway, with multiple proteins being expressed more in the eastern biological replicates, a few being BRI1 and CYCD3. Further comments about these up- and down-regulated pathways are reserved to the Discussion section.

## Discussion

The need to understand what drives local adaptation of forest tree species is perhaps more immediately necessary than row crops due to the time and cost required to rapidly adapt to climate change (Howden et al. 2007). While initial projections of increased temperature and carbon dioxide are expected to increase timber production, sparsity in rainfall and changes in disease or insect pressure may impact any potential gains (Matallana-Ramirez et al. 2021). Furthermore, given that forest plantations typically grow for multiple decades before harvest, as the climate changes, knowing a plantation will thrive even with these changes is essential. Provided the large amount of genetic diversity and known physiological differences among sources of loblolly pine from different geographic regions, there should be interest in understanding what makes these sources different to help mitigate detrimental impacts or take advantage of changing climate. The ability to enable this starts with exploring the fundamental genetic differences that drive local adaption.

To further an understanding of underlying genomic differences between geographic sources of loblolly pine, differential gene expression analysis was conducted between eastern and western loblolly pine families in a common garden experiment. Families were chosen to represent the more extreme natural geographic range of loblolly pine known to have phenotypic differences between them. Due to the limited functional annotation of the loblolly pine transcriptome, we attempted to connect these results back to *Arabidopsis* to gather information on overrepresented gene ontology categories and KEGG pathways in eastern or western sources.

Before performing differential expression, the PCA plot generated from mapped reads displayed apparent differences between the eastern, western, and far-western seed sources. The observation of differences between the eastern and western/far-western sources was not surprising, as multiple studies have displayed the differences between these two provenances (Wells 1983, Schmidtling 2001). However, it was surprising to see the distinction between west and far-western sources. Studies have shown differences between far-western seed sources and western seed sources (Louisiana or Arkansas) at different years of growth and disease resistance (Schmidtling 2001); however, the observation of differences in gene expression patterns in 6-week-old seedlings suggests these differences can be detected even at the seedling stage.

The differential expression results did not display large log2 fold changes between the eastern and western sources; however, given the large number of biological replicates in each category, significance was relatively high for those transcripts identified as differentially expressed. This lack of large changes in gene expression may be due to the absence of treatment, such as water stress, or because young seedlings are inherently expressing the same core genes to establish development, making unique differences between them relatively small. The BLAST results of differentially expressed pine transcripts to *Arabidopsis* cDNA indicated multiple pine transcripts mapping to the same *Arabidopsis* genes in many cases. It is unclear if the multiple mapping is due to high allelic variation within loblolly pine or if transcripts are different members of the same gene family. If the cause is due to allelic variation, it would make sense to consider these multiple mappings the same gene, however if the transcripts are different members of the same gene family it would not. Unfortunately, until a better reference genome assembly is produced from loblolly pine, the only option was to treat them as coming from the same gene.

Given this assumption, the functional annotation of unique *Arabidopsis* genes provided interesting observations. GO categories that were upregulated in the western seed sources included various terms that followed expectations given the known physiological differences, such as response to salt stress, water deprivation, and heat. This finding agrees with loblolly pine studies where treatments are applied (Li et al. 2021). Enriched GO categories for downregulated genes included processes related to phosphorylation, transmembrane transport, and sterol biosynthetic processes. While these terms are not as directly-related to physiological differences as the upregulated categories, fast growth is the most distinguishing factor of eastern versus western seed sources. The genetic mechanisms of how plants can achieve fast growth is complex; however, sterol biosynthetic processes and protein phosphorylation are known to be involved in plant growth and regulation (Schaller 2003, Li and Liu 2021).

When evaluating KEGG pathways, we observed many of the same pathways displaying different up and down-regulated genes. One example is the MAPK signaling pathway, which is involved in various cellular responses within plants to stress and is well documented for its involvement in responses to abiotic pressures (Taj et al. 2010). The observation of several PP2Cs and a SnRK2 gene upregulated for salt and drought tolerance is interesting. Protein Phosphatase 2C (PP2C) has been linked to a response to salt stress in the tree species *Betula platyphylla*, and positively regulates salt and drought tolerance in *Arabidopsis* (Xing et al. 2020, Liu et al. 2012, Bhaskara et al. 2012). Similarly, SNF-1 related protein kinases 2 (SnRK2) have been shown to assist in maintaining plant stability and impacting root system architecture under salt stress (Szymanska et al. 2019, McLoughlin et al. 2012). With respect to the downregulated PYL4 gene, an *Arabidopsis* mutant study found that mutating the PYL4 gene increased interaction with PP2Cs and provided the ability to enhance drought stress (Pizzio et al. 2013).

Another overlapping pathway observed was zeatin biosynthesis. Zeatin is a type of cytokinin, a group of plant hormones known to promote cell division (Wu et al. 2021). We observed downregulation of the *Arabidopsis* response regulator 16 (ARR16) gene within the zeatin biosynthesis pathway. A relatively recent mutation analysis of the ARR16 gene in *Arabidopsis* found it to be a positive regulator for hypocotyl growth in response to blue or white light (Srivastava et al. 2019). The observed upregulated *Arabidopsis* response regulator 1 (ARR1) gene has been found to inhibit root growth in low temperatures (Zhu et al. 2015). These findings may hint at possible ways of moving western sources north to adapt to lower nightly temperatures.

One interesting observation of the pathways that were unique to up or downregulated genes was downregulation of the brassinosteroid pathway. Brassinosteroids are steroid plant hormones involved in many different growth elements, including roots and new green tissue growth (Bajguz et al. 2020). Previous work in apple trees found that application of brassinosteroids enhanced the expression of growth-related genes (Zheng et al. 2019). Among the several genes downregulated in western gulf germplasm, particularly significant were Brassinosteroid-insensitive 1 (BRI1) and D-type cyclin (CYCD3). The overexpression of BRI1 in *Solanum lycopersicum* was found to increase lateral root number, plant height, and hypocotyl length, among many other things (Nie et al. 2017). Studies in poplar (*Populus* sp.) and *Arabidopsis* have associated the overexpression and function of CYCD3 with enhancing new growth (Guan et al. 2021, Dewitte et al. 2007).

The BRITE transcription factor category contained some specific transcription factors upregulated that are known to be involved in stress response. The transcription factors TINY2 and DREB2A are members of the APETALA2/ETHEYLENE RESPONSE FACTOR (AP2/ERF) family, a group of transcription factors found to be important in stress responses (Li et al. 2015). In *Arabidopsis, TINY2* has been found to positively regulate response to drought and inhibit growth driven by the brassinosteroid pathway (Xie et al. 2019). Similarly, the transcription factor DREB2A has been found to be associated with water and heat stress in *Arabidopsis* and maize (Qin et al. 2007, Sakuma et al. 2006). More recently, a case study in rice generated transgenic lines overexpressing both DREB2A and APX, a reactive oxygen species enzyme, and observed increased drought tolerance (Sandhya et al. 2021).

These highlighted findings represent interesting pathways and genes that are related to the known differences among eastern and western seed provenances of *Pinus taeda* in the southern United States. Additionally, they represent fundamental differences in the beginning of seedling development without any treatment or disease pressure applied, showing that there are detectable differences between these two provenances at a young age. Further exploration of the genes identified here is needed prior to using this information in any practical breeding application; however, they represent a starting point in understanding what transcripts may be involved in driving local adaptation of seed sources.

Overall, this experiment contributes to the body of literature on fundamental differences between loblolly pine seed sources. In the future, one could imagine making wide crosses between eastern and western sources to select progeny that have both the fast growth of eastern sources and abiotic tolerance of western sources. Given the length of loblolly pine breeding, this would take time to achieve, but could provide an opportunity to engineer adaptation of families to tolerate the different temperatures and rainfall that climate change will inevitably bring.

## Notes

### Competing Interest Statement

The authors have declared no competing interest.

