## Supplemental Material for "Analysis of Gene Expression Differences Between Eastern and Western Loblolly Pine Seed Sources"

**Supplementary Material**

**Table S1**. List of Family ID's, number of biological replicates, and their corresponding provenance

| Family | # of biological replicates | Group |
|---|---|---|
| LSG111 | 3 | WG |
| TIP1158330 | 3 | TIP |
| TIP138245 | 1 | TIP |
| TIP139150 | 4 | TIP |
| TIP143184 | 1 | TIP |
| TIP181745 | 2 | TIP |
| TIP1857822 | 3 | TIP |
| TIP1932699 | 3 | TIP |
| TIP1973363 | 3 | TIP |
| TIP21206990 | 3 | TIP |
| TIP2751739 | 3 | TIP |
| TIP2818013 | 3 | TIP |
| WG1809 | 3 | WG |
| WG4302 | 3 | WG |
| WG4501 | 3 | WG |
| WG4801 | 3 | WG |
| WG7083 | 4 | WG |
| WG7192 | 3 | WG |
| WG7206 | 3 | WG |
| WG7442 | 3 | WG |
| WG7478 | 1 | WG |
| WG7720 | 3 | WG |

RESTRICTED

**Table S2.** Number of pine transcripts that hit a given *Arabidopsis* gene.

| # of pine transcripts hit | # of *Arabidopsis* Genes |
|---|---|
| 2 | 2247 |
| 3 | 1097 |
| 4 | 596 |
| 5 | 405 |
| 6 | 274 |
| 7 | 223 |
| 8 | 174 |
| 9 | 130 |
| 10 | 118 |
| 10-20 | 507 |
| 21-50 | 292 |
| 51-100 | 74 |
| 101-200 | 29 |
| 201-500 | 13 |
| >500 | 3 |

RESTRICTED
